# Early Life Adversity in Male Mice Sculpts Reward Circuits

**DOI:** 10.1101/2021.07.28.454015

**Authors:** Kara M. Wendel, Annabel K. Short, Brenda P. Noarbe, Elizabeth Haddad, Anton M. Palma, Michael A. Yassa, Tallie Z. Baram, Andre Obenaus

**Affiliations:** Department of Anatomy and Neurobiology, University of California, Irvine School of Medicine, Irvine, CA; Department of Pediatrics, University of California, Irvine School of Medicine, Irvine, CA; Institute for Clinical and Translational Science, University of California, Irvine, Irvine, CA; Department of Neurobiology and Behavior, University of California, Irvine School of Biological Sciences, Irvine, CA

**Keywords:** diffusion tensor imaging, neuroimaging, stress, connectivity, tractography

## Abstract

Early life adversity (ELA) comprises a wide variety of negative experiences during early life and has been linked to cognitive impairments, reduced experiences of pleasure (anhedonia), and other long-term consequences implying that ELA impacts the reward circuitry. In this study, we focused on the projections from the dorsal raphe (DR) to the ventral tegmental area (VTA) and on to the nucleus accumbens (NAcc), an important pathway within the reward circuit. We hypothesized that ELA alters connectivity within the DR-VTA-NAcc pathway, manifested behaviorally as anhedonia in adulthood. We used the limited bedding and nesting model to induce ELA in mice and measured reward-related behaviors in adulthood using the three-chamber social interaction and sucrose preference tests. High resolution *ex vivo* diffusion tensor imaging (DTI) was acquired and processed for regional DTI metrics, including tractography to assess circuit organization. We found brain-wide changes in radial diffusivity (RD) and altered connectivity of the reward circuit in the ELA group. DR-VTA-NAcc circuit tractography and axial diffusivity (AD) along this tract exhibited dispersed organization where AD was increased in the VTA segment. Behaviorally, ELA elicited an anhedonic phenotype in adulthood with decreased direct social approach and time spent with peer but no overt differences in sucrose preference test. Our findings suggest that reward circuits, assessed using DTI, are altered following ELA and that these changes may drive enduring reward deficits.

## 1. Introduction

Circuits begin at the level of the synapse, where integration and propagation of signals amongst axons culminate to drive behavior and cognition; therefore, they are the interpreters of the brain translating electrical signals into behaviors (Tau and Peterson, 2010). The function and structure of these circuits evolve throughout development and are known to be influenced by genetics, environmental experiences, and activity-dependent plasticity (Bale et al., 2010; Bolton et al., 2020; Fox et al., 2010; Hunter and McEwen, 2013; Lucassen et al., 2013; Peña et al., 2017; Suderman et al., 2014; Szyf, 2013; Tau and Peterson, 2010). Circuit maturation requires both genetic and environmental cues, and they take place during sensitive postnatal periods (Birnie et al., 2020; Luby et al., 2020). Because circuits are shaped throughout development and ultimately govern behavior, understanding how developmental perturbations such as early-life adversity (ELA) sculpts circuitry is critical to understanding the long-term impact.

ELA encompasses a multitude of adverse experiences that occur during infancy and childhood which can modulate structure and function of several brain networks and is associated with long-term effects including neuropsychiatric and physical diseases (Chen and Baram, 2016; Elwenspoek et al., 2017; Short and Baram, 2019). Studies have shown that mental development is impaired in infants whose mothers had inconsistent emotional states during and after pregnancy or provided less predictable maternal sensory signals (Davis et al., 2017; Sandman et al., 2012). ELA can be modeled by limited bedding and nesting (LBN), which leads to fragmented maternal care in rodents, resulting in anhedonia-like behaviors among pups: reductions in social play, sucrose, chocolate, and cocaine consumption (Birnie et al., 2020; Bolton et al., 2018a; Bolton et al., 2018b; Goodwill et al., 2019; Molet et al., 2016; Walker et al., 2017). Anhedonia is the inability to feel pleasure in all or almost all activities and is a core feature of reward circuit deficits (Der-Avakian and Markou, 2012).

Previous evidence found that ELA induced long-lasting changes within the VTA via Otx2, a transcription factor critical for dopamine neuron development (Peña et al., 2017). Additionally, ELA in young children has been linked to blunted maturation of VTA functional connectivity (Park et al., 2021). In children, ELA is strongly associated with the development of cognitive and emotional problems later in life, ranging from cognitive deficits to reward deficits and anhedonia (Bremner, 2003; Bremner and Vermetten, 2001; Heim et al., 2008; Nelson et al., 2007; Short and Baram, 2019).

The reward system includes medial prefrontal cortex, the hippocampus, the nucleus accumbens, the amygdala, the ventral tegmental area, and the dorsal raphe amongst other regions (Nestler and Carlezon, 2006). The best characterized portion of the reward circuit is the dopaminergic neurons of the ventral tegmental area (VTA) that project to the nucleus accumbens (NAcc) (Russo and Nestler, 2013). We build on previous studies by investigating the classic reward pathway between the VTA and NAcc but here include the dorsal raphe (DR). Previous studies demonstrated that the DR influences other reward regions. For example, optogenetic stimulation of the DR projection fibers to the VTA excited VTA dopaminergic neurons eliciting NAcc core dopamine release (Qi et al., 2014; Wang et al., 2019). Investigating the DR-VTA-NAcc pathway will uncover the importance of this circuit’s contribution to anhedonia.

The mechanism(s) underlying the long-term outcomes of early life experiences remains unknown. We hypothesized that exposure to ELA alters the maturation of the DR-VTA-NAcc pathway, resulting in enduring changes of connectivity. These may underlie behavioral phenotypes such as anhedonia in adulthood. To test this hypothesis, we induced ELA using the LBN model, performed reward-behavioral tasks, and investigated reward regions and circuits with high resolution diffusion tensor magnetic resonance imaging.

## 2. Methods

### 2.1 Animals and LBN paradigm

All experiments were approved by the Institutional Animal Care Use Committee and carried out according to National Institutes of Health guidelines for experimental animals. All efforts were made to minimize animal suffering and the number of animals used. Primiparous C57Bl/6J female mice were obtained from Jackson Laboratory and were bred in-house. All animals were housed in temperature-controlled, quiet, uncrowded conditions on a 12-h light, 12-h dark schedule (lights on at 0600 h, lights off at 1800 h) with free access to food and water. Females were checked daily for plugs during breeding epochs. At embryonic day 17 pregnant females were singly housed and two cotton nestlets were provided and checked daily for parturition. The day of birth was denoted as postnatal day (PND) 0.

To induce ELA, we employed the LBN paradigm based on Rice et al., 2008. The experimental timeline included: random assignment of dams to control or LBN conditions instituted from PND 2-9, behavioral testing performed from PND 80-100, followed by diffusion tensor imaging (DTI) (Figure 1). Litters that had less than 4 pups were excluded and litters larger than 8 were culled based on sex to maintain a balance between males and females. Control dams were placed into standard cages containing a normal amount of bedding and one cotton nestlet. In ELA cages the bedding was reduced to scarcely cover the cage bottom, a fine gauge aluminum mesh was placed above the cage floor, and half of a cotton nestlet was provided (50% reduction). From PND 2-9, cages were left undisturbed during which maternal observations were performed via video recordings. On PND 10, all cages reverted to standard cages with normal bedding and nesting.

**Figure 1.**
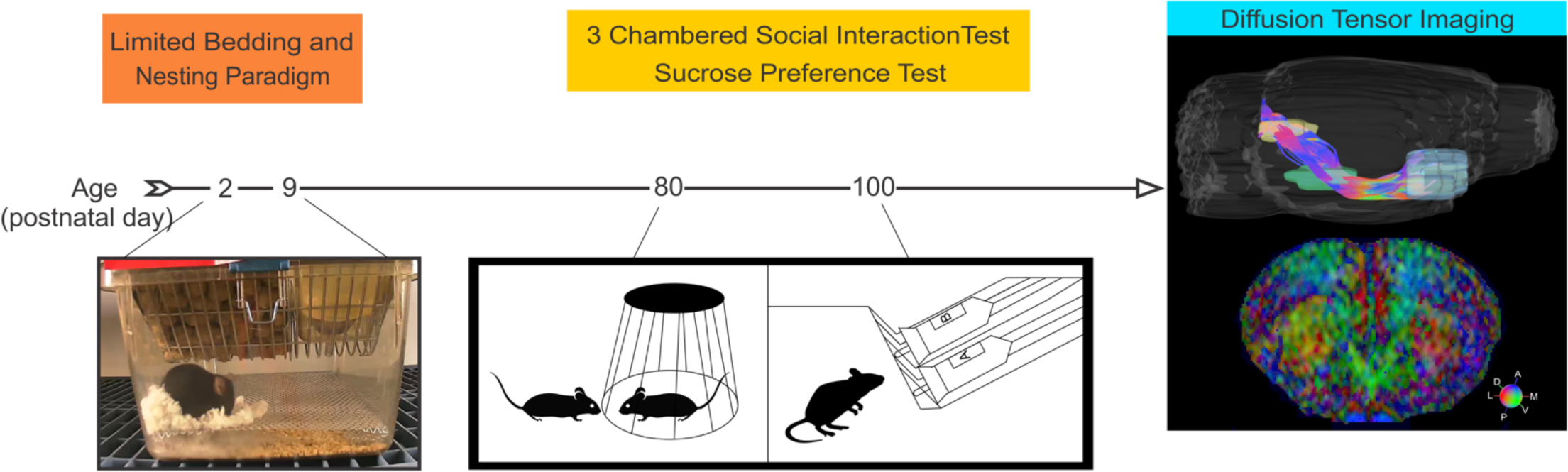
Experimental Timeline. Schematic for assessment of reward-related behavioral tasks and diffusion tensor imaging (DTI) following ELA.

### 2.2 Anhedonia testing in adulthood

Behavioral testing was performed PND 80-100 and two reward-related tasks to assess anhedonia were performed: three-chambered social interaction and sucrose preference tests. Male mice (n=18 controls, n=25 ELA mice from 12 litters) were placed into a three chambered arena, in which they spent 10 minutes habituating to the center chamber, then to all three empty chambers. Mice then had 10 minutes to interact with a novel object (empty cup) or a stranger mouse placed inside a cup (Yang et al., 2011). Time spent with the novel mouse versus the empty cup was quantified as a measure of pleasure from social interaction. After the three-chambered social interaction test, the sucrose preference test was performed for 3 days. Bottles of both 1% sucrose and drinking water were given *ad libitum* and measured each day of a three-day testing period for each animal. Bottles were switched from the left to the right and vice versa each day in order to prevent and identify side preference. Sucrose preference was quantified as the percentage of sucrose consumed out of total liquid consumption for each day (sucrose divided by the sum of sucrose and water). Experimenters were blinded to group designation during behavioral testing and analysis. Animals that did not perform the tasks such as those with side preference or those that only consumed sucrose were excluded.

### 2.3 MRI acquisition

After behavioral testing, mice were sacrificed via transcardial perfusion with phosphate buffered saline (PBS) and 4% paraformaldehyde (PFA). The brain was kept in the cranium and then post-fixed in 4% PFA overnight followed by three changes of 1x PBS over 3 days, after which they were stored in PBS with sodium azide (0.02%) until imaging was completed. Brains (n=13 controls, n=21 ELA) then underwent *ex vivo* diffusion tensor imaging (DTI) and T2-weighed imaging (T2WI) using a 9.4 T Bruker Avance imager (Bruker Biospin, Billercia, MA). All data were collected with the following parameters: 1.5-cm field of view, 0.5-mm slice thickness, and a 128x128 acquisition zero filled to a final 256x256 matrix. DTI parameters were: repetition time (TR)/echo time (TE) = 8000 msec/35.66 msec, 30 isolinear directions, b = 3000 mT/m, with 5 Bo images acquired prior to weighted images. T2WI parameters included TR/TE = 4000/10 msec with 10 equally spaced echoes. Total imaging time was 1 h, 52min.

### 2.4 MRI analysis

Experimenters were blinded to group designation during MRI analysis. MRI scans were eddy corrected and the cranium was removed using MATLAB. FMRIB’s Diffusion Toolbox from FMRIB’s Software Library (FSL) was used to generate parametric DTI maps, in which a diffusion tensor model was fit at each voxel.

Each animal’s T2 and b0 images were linearly registered to create a T2/b0 map. The Australian Mouse Brain Atlas Consortium (AMBMC) atlas was then non-linearly registered to each individual animal’s T2/b0 map and regional labels were applied with Advanced Normalization Tools (ANTs). Regional statistics for fractional anisotropy (FA), mean diffusivity (MD), axial diffusivity (AD), and radial diffusivity (RD) were extracted for 83 regions (41 bilateral regions, 1 midline region). FA reports the asymmetry of water diffusion (FA = 0, isotropic diffusion; FA = 1, anisotropic diffusion), MD reports the average magnitude of water diffusion in the tissue of interest, AD reflects diffusion of water parallel to axons, and RD manifests water mobility perpendicular to axons (Obenaus and Jacobs, 2007; Song et al., 2003). Percent change was computed from the group averages to the controls for each region and each metric.

Scans were then reconstructed in DSI studio using a diffusion sampling length ratio of 0.75 (April 11, 2018 build; http://dsi-studio.labsolver.org) and AMBMC region designations were used to generate tractography. The DR was used as the seed and the VTA and NAcc were used as regions of interest.

Deterministic tractography with FIB Autocorrect was performed using the following global parameters: angular threshold=60, step size=0.06, smoothing=0.60. FIB Autocorrect is a method that tests if each fiber orientation forms a coherent track that connects to the neighboring voxel, and slightly corrects its orientation accordingly. Tract statistics were exported, and tract data were exported for the FA, MD, AD, and RD along the fiber orientation. The DR-VTA-NAcc tract was split into five segments: DR, S1 (tract between DR and VTA), VTA, S2 (tract between VTA and NAcc), and NAcc in order to localize changes. Segment averages were computed by identifying the center of the segment and sampling 25% above and below the middle point and then averaged to determine the FA, MD, AD, and RD average of the tract as it passes through and between regions.

In addition, DTI scans were reconstructed in DSI Studio with a diffusion sampling length ratio of 0.85.

Whole brain seeding was performed and the reward regions from the atlas registration were selected to generate a connectivity matrix of the reward circuits. The connectograms of each animal were then averaged within the group using Microsoft Excel. Group averaged data was then plotted using Circos http://circos.ca/, (Krzywinski et al., 2009). For the reward connectograms only the 4^th^ quartile was selected in Circos (Figure 3A), and for DR-VTA-NAcc connectograms all quartiles were included (Figure 3D). The colors correspond to each region and the thickness of the ribbons correspond to the strength of connectivity.

### 2.5 Normalizing tract lengths from tractography

To normalize tract length for each animal, the tract was split into segments (see Figure 3C) and the length of each segment was determined based on the animal with the longest segments. Data were then spaced in order to standardize the tract length across animals and Real Statistics Resource Pack (https://www.real-statistics.com) was used to perform imputation for missing data. The data was imputed by group and by segment to account for variation within different segments of each tract. A rolling average of the two positions before and after was applied to each tract to minimize noise. One animal who was an outlier in every DTI metric was normalized to the mean, based on the percent change of each metric.

### 2.6 Statistical analysis

All statistics were computed using Graphpad Prism 8 with 1.5 interquartile range-based outlier analysis using Microsoft Excel. Kolmogorov-Smirnov normality tests were run with the normalized means to determine if parametric testing should be used. T-tests were used to assess the differences between control and ELA groups unless otherwise stated. When groups failed normality tests, Mann-Whitney tests were used in lieu of t-tests. Using the VTA RD, animals that were greater than 2.5 standard deviations above the control mean were classified as ELA-high RD and the remainder of the ELA animals were classified as ELA-low RD. One-way ANOVA was used to assess differences between control, ELA-high RD, and ELA-low RD groups and two-way repeated measures ANOVA was used to analyze along tract metrics. Post-hoc tests were performed with Tukey’s testing. Correlative statistics were determined using a linear regression testing to explore the relationships between sucrose preference data and tract metrics. Data are presented with mean and standard error of the mean (SEM) and a p < 0.05 was used to assess statistical significance.

## 3. Results

### 3.1 Brain wide regional changes in radial diffusivity suggest widespread alterations after ELA

Assessment of brain wide diffusion-derived metrics was performed for 41 bilateral regions in the mouse brain. A broad survey of the resultant regional diffusion data (164 data points) in ELA mice compared to controls found that axial diffusivity (AD) and radial diffusivity (RD) revealed widespread diffusion changes after ELA (Figure 2). Remarkably, global AD increases were observed in the ELA mice whilst every region of the ELA group had increased RD. On average, relative to controls, the percent change in RD was 18.5%, whereas the average percent change in AD was a more modest 2.0%. The number of regions with increased or decreased AD and RD are illustrated in pie charts with virtually all regions exhibiting positive percent change in RD (Figure 2).

**Figure 2:**
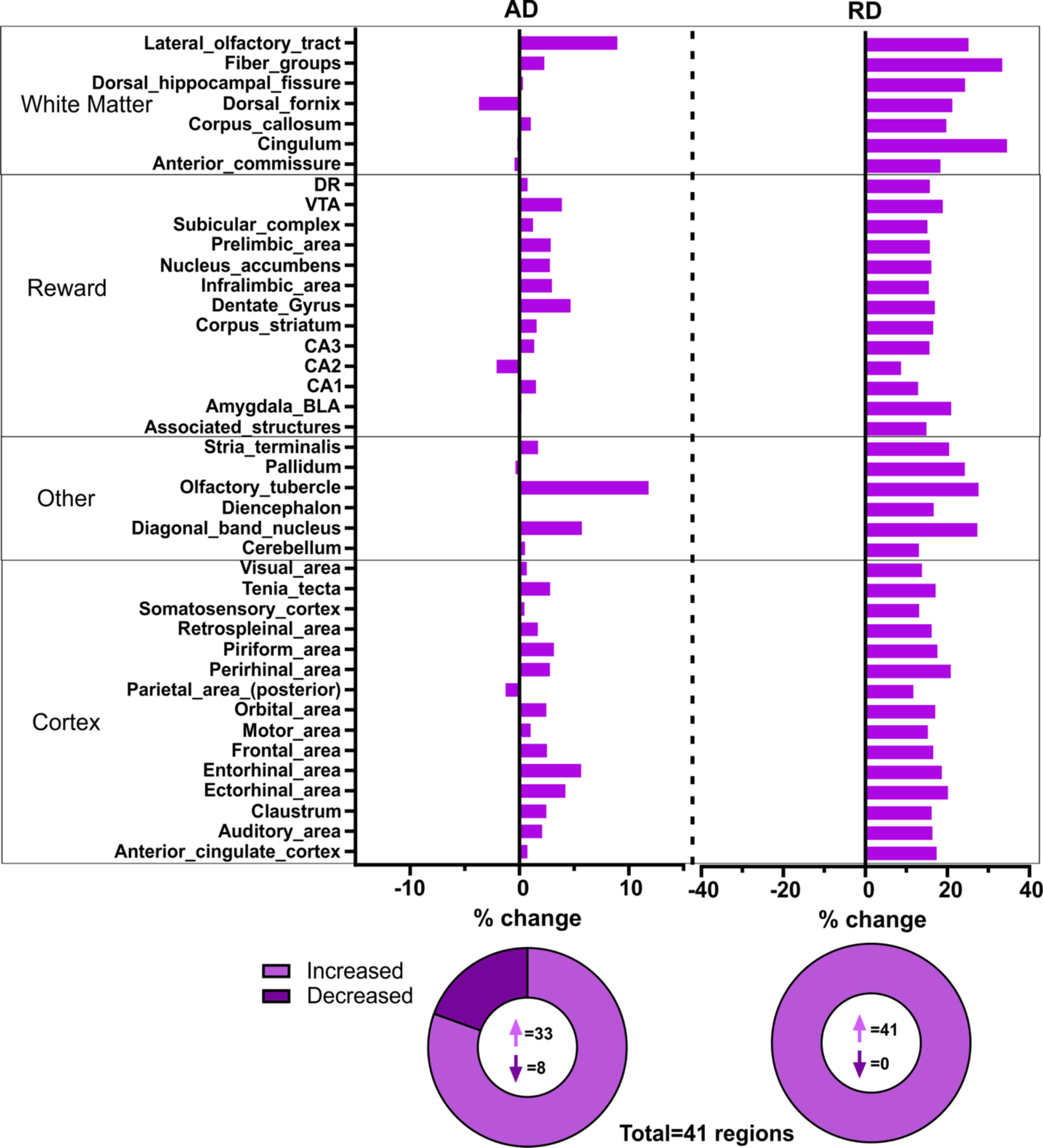
Regional brain axial diffusivity (AD) and radial diffusivity (RD) after ELA. Percent change in AD and RD of the ELA mice compared to controls revealed that many regions were impacted after ELA. There was a strikingly increase in AD across almost all brain regions and a dramatic increase in RD across all regions after ELA. Pie charts show the distribution of brain regions with increased versus decreased percent change for AD and RD in the ELA group. Majority of brain regions exhibited increased percent change in AD and all regions had increased RD.

FA and MD of the ELA group also exhibited global differences compared to controls (Supplemental Figure 1). Decreases in FA were identified in ELA mice (-7.2%) and increased MD (3.8%) was found in the most regions. AD and RD both contribute to MD, therefore MD is not as illuminating as AD and RD alone. As such, in the current study RD was the best predictor of brain-wide changes following ELA.

### 3.2 Reward circuit connectivity is increased, and organization is dispersed in ELA mice

Based on the observed global regional changes, we next assessed reward circuit connectograms in control and ELA mice. Connectograms illustrated altered connectivity between anhedonic regions (Figure 3A) with stronger connectivity from the DR to VTA in ELA mice compared to controls. There was also increased left VTA connectivity in ELA group compared to control mice. The connectivity of the right basolateral amygdala (BLA) was also increased in the ELA group. Since the DR and VTA exhibited strong changes in connectivity, the DR-VTA-NAcc projection was then mapped using tractography to visualize altered tract organization in ELA mice (Figure 3B). For quantitative analyses, the tract was mapped into five distinct components, the tracts in the DR, VTA, and NAcc, along with tracts connecting the DR/VTA (S1), and tracts between VTA/NAcc (S2) (Figure 3). As shown in Figure 3B (arrows), the tract integrity was dispersed within the S2 and NAcc of ELA mice compared to control mice. Connectomes of only DR-VTA-NAcc projections from control and ELA mice further emphasized altered connectivity (Figure 3D). The connectivity of the ELA group was decreased between the left and right NAcc but increased between the left VTA and NAcc in the ELA group compared to control mice (* in Figure 3D). This was further emphasized by altered connectograms of just DR, VTA, and NAcc regional connectivity (Supplemental Figure 2). The ELA group exhibited stronger DR connectivity to the left VTA, decreased connectivity between DR and the right NAcc, and left VTA (*). The connectivity of the left VTA was increased to the NAcc, but exhibited connectivity to the DR in the ELA group. The left NAcc exhibited increased connectivity to the left VTA but decreased connectivity to the right NAcc in ELA mice. The connectivity and organization of the DR-VTA-NAcc tract demonstrate that the reward circuit is modified after ELA.

**Figure 3:**
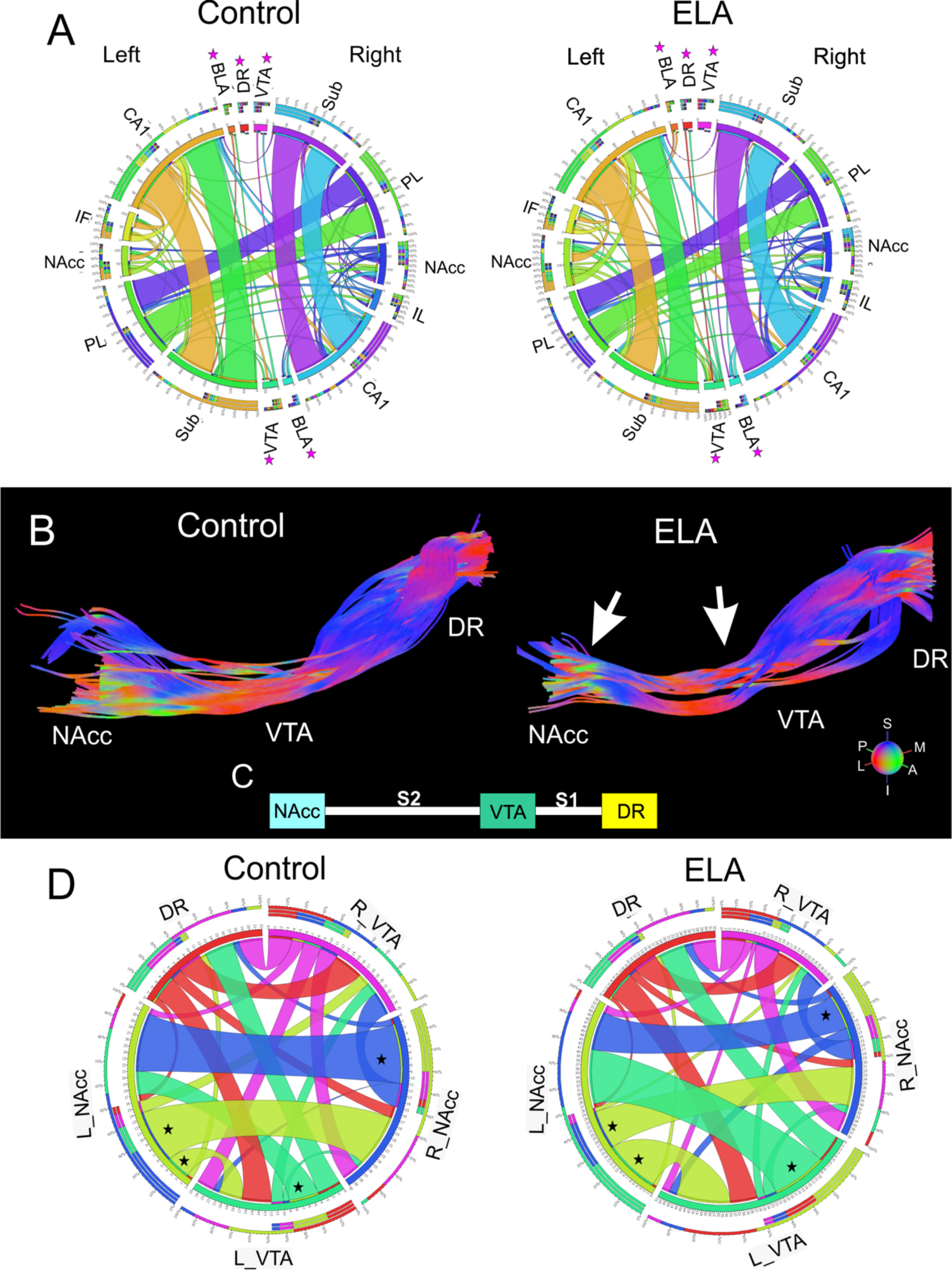
Organization of reward circuits were altered following ELA. A) Diffusion tensor imaging (DTI) reward circuit connectograms of control and ELA mice illustrated altered connectivity between anhedonic regions. Altered connectivity between control and ELA groups was particularly evident in BLA, DR and VTA regions (*). B) DTI tractography mapped from the DR through the left VTA and terminating in the left NAcc of control and ELA mice. As can be visualized, ELA mice exhibited increased tract dispersion between VTA and NAcc. C) Block diagram represents the different tract segments where S1 (segment 1) and S2 (segment 2) are the tract portions between the nuclei (DR, VTA, NAcc). D) DTI connectograms focusing on the DR-VTA-NAcc circuit in control and ELA mice further highlighted altered connectivity. The connectivity of the ELA group exhibited different connectivity between the VTA and NAcc (*).

### 3.3 Regional DR, VTA, and NAcc diffusion metrics

When regional brain diffusion metrics were assessed in DR, VTA, and NAcc, RD exhibited significant alterations (Figure 4). No differences were found between the ELA and control groups in AD for the DR, VTA, or NAcc (Figure 4A). In contrast, RD exhibited significant changes in the DR, VTA, and NAcc (Figure 4B). A significant increase in DR RD (t=2.632, df=31, p=0.013), in VTA RD (Mann-Whitney, p=0.0299) and in NAcc RD (t=2.712, df=31, p=0.011) were found within the ELA group relative to controls. These findings hint that in ELA mice the DR, VTA, and NAcc are sensitive to RD changes.

**Figure 4:**
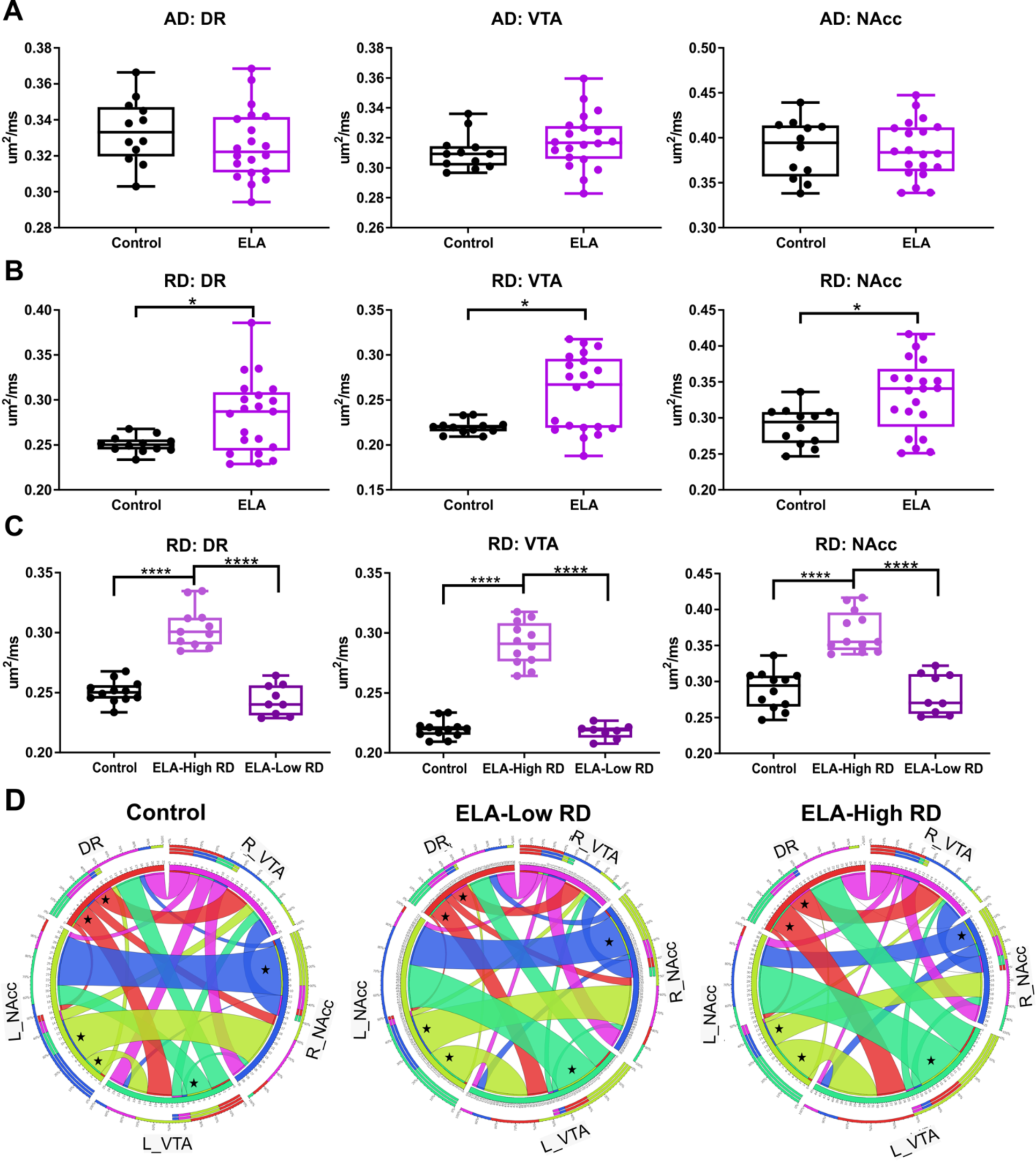
DTI metrics within the DR, VTA, and NAcc showed changes in radial diffusion. A) Regional axial diffusivity (AD) exhibited no changes in AD within the DR, VTA or NAcc. B) Radial diffusivity (RD) demonstrated a significant increase in the RD within the DR, VTA, and NAcc of the ELA group. There was the appearance of subgroups in the RD within the VTA. C) Radial diffusivity (RD) was divided into ELA-high RD and low RD which revealed significantly increased RD in the ELA-high group compared to controls and ELA- low RD groups within the DR, VTA, and NAcc. D) DTI connectograms for the DR-VTA-NAcc circuit of control, ELA-low RD, and ELA-high RD mice highlighted altered connectivity where ELA-low RD animals more closely mirrored the connectivity of controls. The connectivity of the ELA-high RD group exhibited different connectivity between the VTA and NAcc, left and right NAcc connections, and connectivity of the DR to the VTA and NAcc (*). (* *p<*0.05, ** *p<*0.01, *** *p<*0.001, **** *p<*0.0001).

The VTA RD of the ELA group failed normality tests suggesting the possibility of two groups within the ELA animals. Animals were classified as ELA-high RD or low RD based on the RD within the VTA (Figure 4C). One-way ANOVA revealed a significant difference in RD of the DR (F (2, 29) = 65.9, p<0.0001), the VTA (F (2,29) = 127, p<0.0001), and the NAcc (F (2, 30) = 33.71, p<0.0001) between groups. Post hoc testing confirmed that there was a significant increase in the RD of the DR (p<0.0001) in the ELA high-RD group compared to the control and ELA-low RD groups. There was also a significant increase in the RD of the VTA (p<0.0001) and of the NAcc (p<0.0001) in the ELA-high RD group compared to the control mice and the ELA-low RD groups. DTI connectograms of the ELA-low RD and high RD groups further confirmed the exacerbated connectivity in the ELA-high RD group (Figure 4D). The connectivity of the ELA-low RD group resembled that of the controls, but the connectivity within the ELA-high RD group was decreased between the left and right NAcc with increased connectivity between the left VTA and NAcc and between the DR and the left and right VTA (*). Thus, regional RD metrics of the DR-VTA-NAcc suggest a modified circuit after ELA.

### 3.4 Along the tract metrics confirm altered organization of DR-VTA-NAcc connectivity

To further assess modifications to DR-VTA-NAcc projection DTI metrics were sampled along the tract (Figure 5). Two-way ANOVA of AD along the DR-VTA-NAcc tract revealed a significant interaction of region and group (F (315, 5985) = 1.848, p<0.0001), and significant effect of region F (4.458, 84.69) = 3.203, p=0.014) (Figure 5A). In ELA mice, a lower AD in the DR was maintained as it coursed to the VTA, whereupon entry to the VTA, the AD was then increased relative to controls. Then AD modestly increased after exiting the VTA as it projected to the NAcc, with a sharp increase in AD as it entered the NAcc. These within tract AD profiles can be readily observed when plotted as a percent difference from controls highlighting the large deviations from controls in the ELA group (Figure 5B). The differences in AD suggest that tract organization within the VTA and NAcc segments is modified after ELA compared to control mice. Increased AD can reflect complex set of interactions evolving from multiple biological factors including increased axonal fiber coherence.

**Figure 5.**
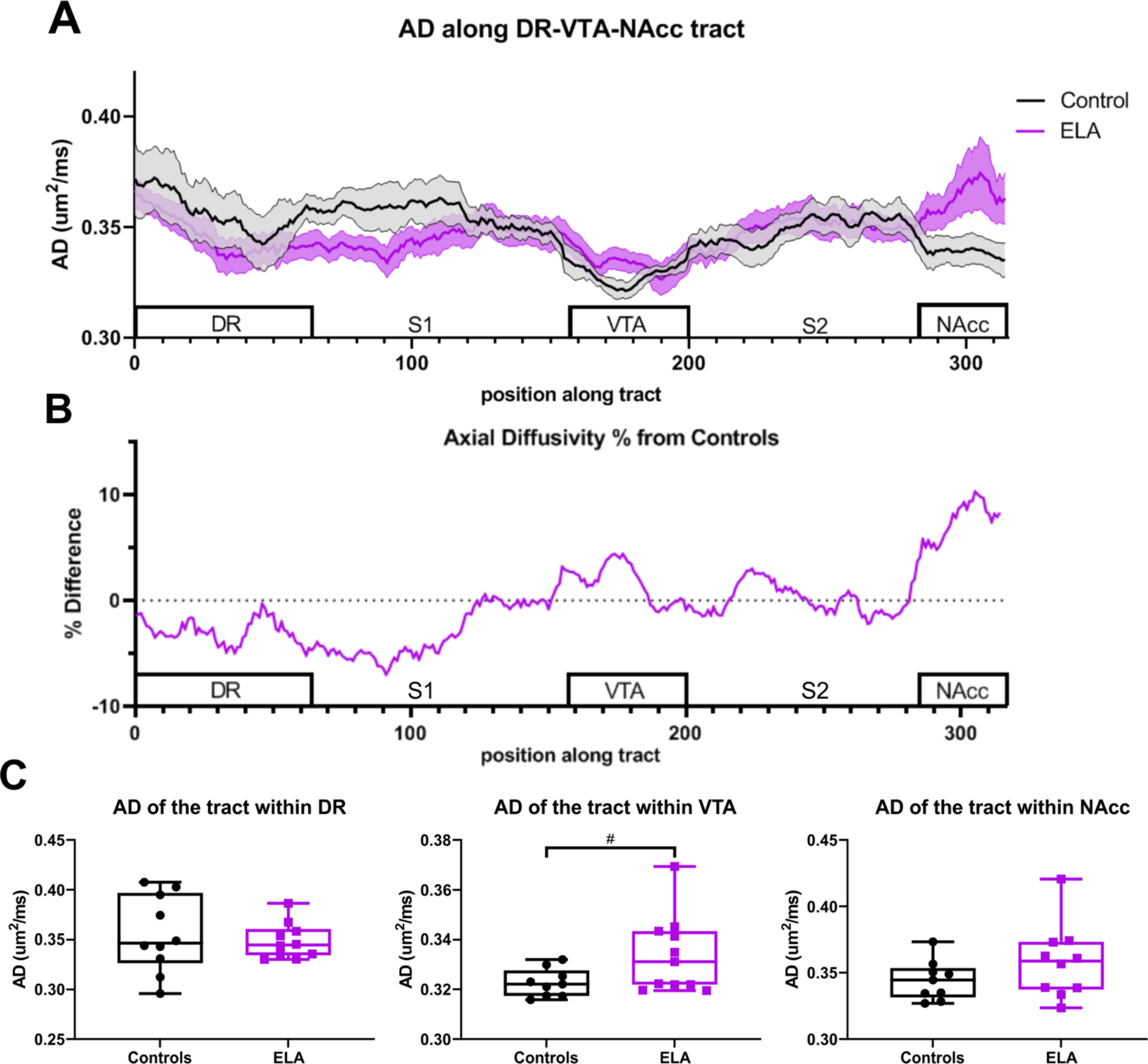
Axial Diffusivity (AD) showed changes within DR-VTA-NAcc projection. A) AD along the tract from DR to VTA to NAcc diverged between the two groups. The DR AD of ELA mice was low, with continued decreases in its connection to and within the VTA. There was a slow but progressive AD increase after exiting the VTA as it projected to the NAcc, with a sharp increase in AD in the NAcc. B) The percent difference of the along the tract relative to controls further illustrates these altered tracts in ELA mice. C) Tract AD within each region revealed a trending increase in AD of the tract within the VTA of the ELA group compared to controls. (# *p<*0.06, * *p<*0.05).

Along the tract MD also exhibited a significant interaction between region and group (F (315, 5985) = 2.562, p<0.0001) and an effect of region (F (3.974, 75.51) = 11.25, p<0.0001). Similar alterations were observed in along the tract RD metrics with significant interaction of region and group (F (315, 5985) = 2.348, p<0.0001) and effect of region (F (4.099, 77.88) = 17.96, p<0.0001). Unlike the tumultuous AD along the DR- VTA-NAcc tract, both MD and RD along the tract of the ELA group were decreased in the DR and remained relatively unchanged across the S1 and VTA, but the S2 it progressively increased peaking in the NAcc. The sharp increase in MD and RD in addition to the AD upon entry into the NAcc of ELA mice suggest that the tract fibers within the NAcc are susceptible to the effects of ELA.

We also averaged the AD within each segment of the DR-VTA-NAcc tract to determine if there was a particular regional tract segment that was more sensitive. No significant changes in AD, within the DR or within the NAcc were found between groups (Figure 5C). However, a trending increase in AD within the VTA was found in ELA mice compared to controls (t=2.032, df=18, *p*=0.057) (Figure 5C). A significant decrease was found in MD within the tract of the DR of the ELA group compared to controls (t=2.246, df=16, p=0.039). The tract within the NAcc also had a trending increase in MD in the ELA group (t=1.891, df=18, p=0.075).

Conversely, RD of the tract segments did not exhibit significant changes. Therefore, the organization of the tract as it passes through the VTA and NAcc are modified by ELA as reported by AD and MD.

### 3.5 Behavior reveals sociability deficits after ELA

Anhedonic behavioral data revealed the emergence of a unique phenotype following ELA. In the three-chamber social interaction, ELA mice exhibited sociability deficits (Figure 6A-C) where ELA mice had significant reductions in direct social approach (t=2.687, df=37, *p*=0.011) and average time spent with peer (t=3.254, df=40, *p*=0.002) when compared to controls (Figure 6A, 6B). No significant differences between groups were found in the average time spent with the object, consistent with the emergence of sociability deficits in ELA mice (Figure 6C). Sucrose preference was also used to monitor anhedonic behaviors and the ELA group was not significantly different from controls across the three days. The three-chamber social interaction data suggest sociability deficits within ELA mice.

**Figure 6:**
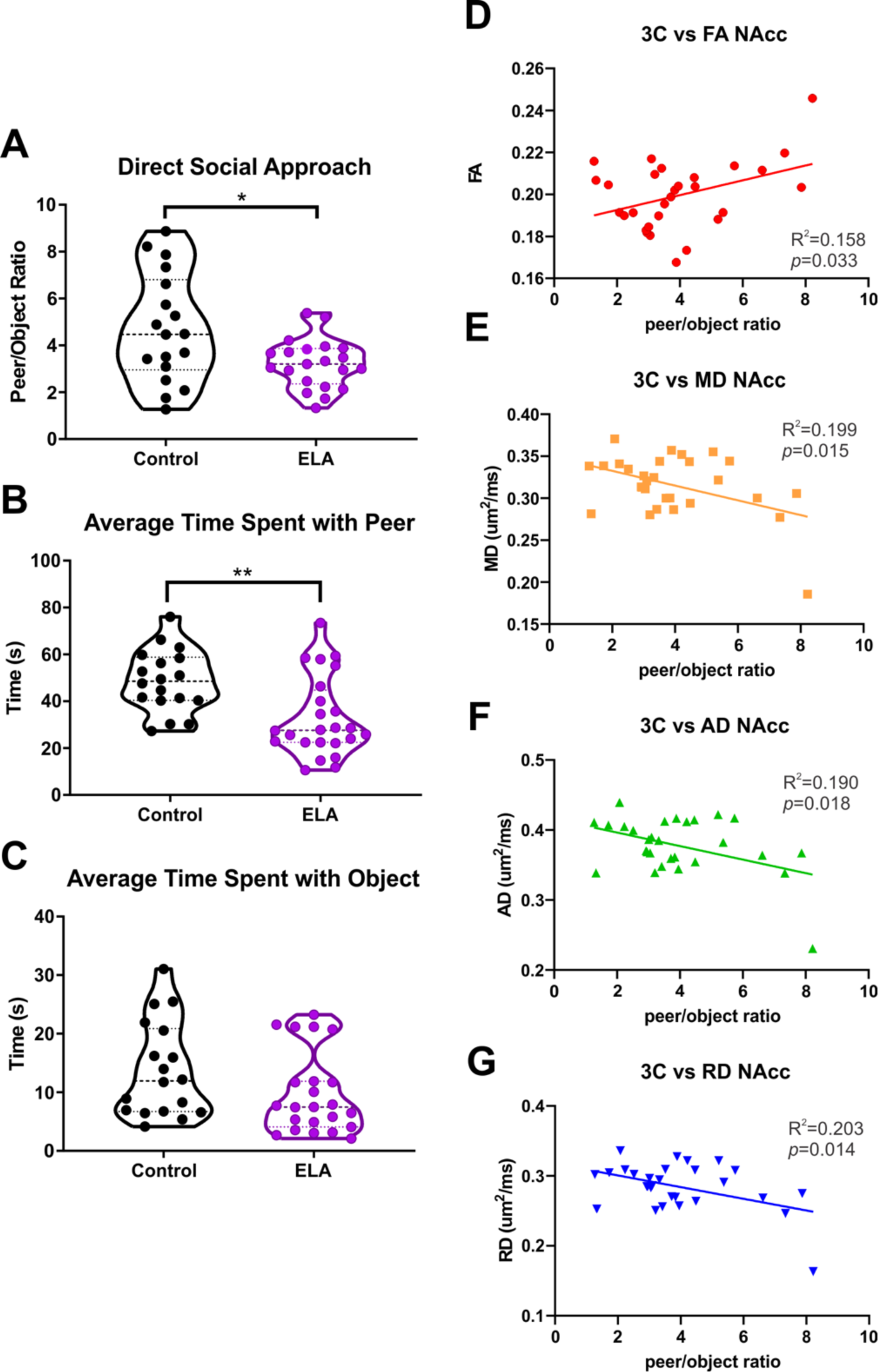
Social reward deficits emerged with the three-chambered social interaction test and correlated with DTI regional metrics in the NAcc. A) Direct social approach highlighted that ELA mice had a significant decrease in peer/object ratio consistent with an anhedonic phenotype after ELA. B) ELA mice revealed a significant decrease in the amount of time spent with a peer mouse. C) No differences between groups in the time spent with the object were observed. D) Peer/object ratio was significantly correlated with the FA of the NAcc. E) Peer/object ratio was negatively correlated with MD of the NAcc. F) Similarly, peer/object ratio was negatively correlated with AD of the NAcc. G) Peer/object ratio was also negatively correlated with RD of the NAcc. (* *p<*0.05, ** *p<*0.01).

### 3.6 Behavioral metrics correlate with diffusion metrics in the NAcc

The behavioral and tissue diffusion changes that we observed were further explored to examine potential inter-relationships. The direct social approach was significantly correlated with NAcc metrics (Figure 6D-G): FA (*p*=0.033; R^2^=0.158), MD (*p*=0.015; R^2^=0.199), AD (*p*=0.018; R^2^=0.190), and RD (*p*=0.014; R^2^=0.203). These correlative relationships between direct social approach and the DTI metrics of the NAcc imply that sociability deficits could be related to alterations within the NAcc. Moreover, these correlations advocate that the NAcc is the most predictive of reward behavioral changes.

## 4. Discussion

The principal findings of this study are that organization and connectivity within the DR-VTA-NAcc projection is enduringly altered following ELA, and these alterations associate with and might contribute to sociability deficits. Using behavioral and DTI indices we found 1) brain wide alterations in radial diffusivity (RD) that arose after ELA, 2) reward circuit connectivity was increased in ELA mice, 3) RD reflected regional changes after ELA in the DR, VTA and NAcc and was able to distinguish ELA animals with a severe phenotype, 4) along the tract DTI metrics unveiled altered organization within DR-VTA-NAcc projection, most prominently within VTA and NAcc, 5) decreased peer/object ratio indicating sociability deficits following ELA, and 6) three-chamber social interaction was well correlated to DTI metrics in the NAcc. Thus, diffusion MRI (dMRI) can identify the ELA phenotype that relates to behavioral outcomes.

### 4.1 Brain wide changes in DTI highlight impact of ELA

To probe the brain on a global scale, we utilized DTI to map regional and connectivity alterations.

Clinically relevant DTI magnetic resonance imaging (MRI) can assess the underlying microstructure of tissue based on water diffusion in the brain (Obenaus and Jacobs, 2007). DTI metrics include FA that reports the asymmetry of water diffusion, MD details the average magnitude of water diffusion, AD reflects diffusion of water parallel to axons and is the largest diffusion vector, while RD manifests water mobility perpendicular to axons or to the largest directional diffusion vector (Song et al., 2003). We observed regional changes in RD and AD following ELA, with overt global increases in RD. Additionally, FA and MD also had large alterations in the ELA group. We found that brain wide changes in RD were largest in gray and white matter integrity following ELA.

FA is the most widely reported DTI metric. Wide-spread gray and white matter integrity changes have been shown in previous studies following ELA with increased FA in the mouse sensory cortex, hypothalamus, amygdala and decreased FA in the corpus callosum and stria medularis following ELA (White et al., 2020). We have previously also reported increased in FA in the CA1 of adolescent rats after ELA (Molet et al., 2016). In our current study, we found a widespread FA decreases following ELA, consistent with findings from non-human primates where decreased FA of the middle longitudinal fasciculus and the inferior longitudinal fasciculus were reported (Howell et al., 2019). In adolescent maltreated rhesus monkeys, significant reductions in FA and increases in RD in corpus callosum, occipital white matter, and external medullary were found which is consistent with our findings (Howell et al., 2013). While we observed reductions in FA, RD was the DTI metric that exhibited the greatest magnitude of regional changes.

Widespread RD increases were found throughout the brain after ELA in our study. This may predict an increased vulnerability to anhedonia after ELA. In a study of depression in adolescents increased anhedonia severity was associated with greater RD in the anterior limb of internal capsule and posterior cingulum both of which are involved in emotion and reward processing (Henderson et al., 2013). In adult depression, a significant positive correlation between anhedonia and increased RD in the anterior limb of the internal capsule was found (Dillon et al., 2018). Therefore, the widespread increases in RD may be indicative of an anhedonic phenotype or vulnerability to reward deficits such as anhedonia.

### 4.2 Reward circuit connectivity is increased after ELA

The broad regional global changes in our ELA mice prompted us to refine our focus on the reward circuit, specifically the DR-VTA-NAcc connectivity. DTI connectograms found that the ELA group exhibited decreased connectivity between the left and right NAcc but increased connectivity between the left VTA and NAcc compared to control mice. This altered connectivity of the DR-VTA-NAcc projection is consistent with ELA-induced modifications of the reward circuit. There are few reports on structural connectivity of the reward circuit after ELA. White and colleagues reported reduced global network efficiency, which is the ability of a network to propagate parallel information, and increased small-worldness indicative of a premature network after ELA (White et al., 2020). Our results similarly find altered hemispheric connectivity. In a study of children, increased generalized FA of the uncinate fasciculus, connecting amygdala to the orbitofrontal cortex, was associated with high maternal unpredictability. (Granger et al., 2021). While these global analyses examined multiple brain circuits, including reward circuits, the findings are consistent with abnormal structural maturation after ELA. In Parkinson’s disease, in which anhedonia is a prominent symptom, connectivity of the reward circuit identified increased AD in connectivity from the substantia nigra (another dopamine hub) to the posterior putamen and globus pallidum, reduced FA in the basal ganglia-motor cortex pathway, and reduced orientation dispersion index in the amygdala-accumbens-pallidum circuit (Guo et al., 2020). The increased AD and reduced FA is similar to our global increases in AD and reductions in FA of the ELA group. Children who experienced ELA exhibited distinct associations of FA within white matter microstructure in tracts linking the striatum and PFC (Dennison et al., 2019). Overall, these data point to neuropathology linked to anhedonia that is manifested as altered connectivity of reward circuits.

Previous clinical evidence from individuals with moderate to severe childhood trauma reported elevated physical, social, and anticipatory anhedonia with decreased left NAcc-right orbital frontal cortex functional connectivity (Fan et al., 2021). Children and mice exhibited altered amygdala development and heightened activity following ELA (Malter Cohen et al., 2013). Additionally in children, a more mature functional connectivity between the amygdala and PFC was displayed following ELA (Gee et al., 2013). Amygdala hyperconnectivity with the hippocampus in ELA mice is also congruent with the observed increased connectivity of the basolateral amygdala within the ELA mice of this study (Johnson et al., 2018). Increased connectivity of the reward circuit has been suggested to be an adaptive response to ELA (Herzberg and Gunnar, 2020) and is consistent with extensive connectivity alterations within the reward system of ELA mice compared to controls. In sum, this suggests blunted hedonic responses via increased connectivity within the reward system as a result of childhood trauma or ELA.

### 4.3 Regional RD alterations in DR, VTA, and NAcc of ELA mice

Regional diffusion metrics of the DR-VTA-NAcc projection found significant increases in RD in ELA mice in all three regions. RD was used to dichotomize those mice severely affected by ELA as observed in both regional metrics and increased connectivity between the DR and VTA and between the VTA and NAcc. These findings confirm that the DR, VTA, and NAcc are ELA sensitive regions. Increased AD/RD can reflect reduced cellular density (Qiu et al., 2008). The presence of crossing fibers, inflammation, and progressive injury are part of different cellular responses that result in DTI metric anomalies (Winklewski et al., 2018). It is difficult to determine what underlying pathology or cellularity changes drove our regional RD alterations within the DR, VTA, and NAcc; to address this gap we further explored the DR-VTA-NAcc projection with tractography.

### 4.4 Organization of the DR-VTA-NAcc projection is altered

Prompted by the increased connectivity in the DR and VTA, we next examined tractography of the DR- VTA-NAcc projection. ELA mice exhibited more dispersed fibers within the VTA and NAcc. When assessing the diffusion metrics along the DR-VTA-NAcc tract, we observed a distinct AD pattern along the tract of ELA compared to control mice, with the largest changes centralized to the VTA and NAcc. These findings further confirm that the tract organization as it passes through the VTA and NAcc are sculpted by ELA.

We and others have previously shown altered connectivity within reward circuits following ELA, with increased streamlines (from tractography) and disorganized connectivity between the amygdala and PFC in ELA rats (Bolton et al., 2018a) and confirmed by increased streamlines between amygdala and hippocampus (White et al., 2020). In conjunction with evidence of other altered reward circuits, our study supports ELA-induced modifications to circuits within the reward system. To the best of our knowledge this is the first study to have sampled and quantified diffusion metrics along the tract to assess region-specific microstructure following ELA. The increased VTA AD suggests a more cohesive fiber organization; in which ELA mice have cohesive axonal bundles as the tract runs through the VTA, in contrast to less organized fibers within the control mice (Dubois et al., 2008). As noted above, these changes in tract characteristics may be due to accelerated maturational signals. Increased AD correlated with an increase in fiber coherence in Parkinson’s disease (Guo et al., 2020). Other studies support that an increased AD reflects smaller axonal diameters within a tract, providing fibers more uniformity due to higher interaxonal space (Kumar et al., 2012; Suzuki et al., 2003). The structural and functional activity of the VTA has also been reported to be altered, where increased excitability of presumed dopaminergic neurons in the medial VTA was reported, with increased spine head diameter, a correlate of synaptic activity (Spyrka et al., 2020). Together with evidence of increased dopaminergic neuronal density in the VTA (Kapor et al., 2020) and increased AD of the tract within the VTA of our ELA mice suggest a delayed pruning of the connectivity of the DR-VTA-NAcc projection resulting in increased excitability, increased fiber coherence, and tract density within the VTA of ELA animals. Thus, AD is predictive of circuit organizational changes, whereas RD reveals changes within the regions as a whole.

The increased MD of the tract within the NAcc of the ELA group further confirmed that both the VTA and NAcc are ELA sensitive regions. Increased MD can reflect a decrease in white matter integrity (Solowij et al., 2017). Alternatively it may be related to hypertrophy of medium spiny neurons within the NAcc, which are associated with stress-induced anhedonia (Bessa et al., 2013). The changes in MD of the tract within the NAcc of the ELA animals likely indicate hypertrophy or reduced axonal packing based on the organization of tracts in the NAcc.

### 4.5 Sociability deficits arise after ELA

ELA is also associated with increased vulnerability to depression, anhedonia, drug use and addiction, as well as cognitive deficits (Heim et al., 2008; Nelson et al., 2007; O’Connor et al., 2015). Anhedonia is the inability to feel pleasure, it is often a prominent symptom of many neuropsychiatric disorders (Hoflich et al., 2019) and is recognized as a critical Research Domain Criterion by the National Institute of Mental Health.

Anhedonia can encompass other reward-related deficits such as motivation, valuation, and decision-making (Der-Avakian and Markou, 2012). A prominent subdomain of anhedonia is sociability deficits as highlighted by tests like the Anticipatory and Consummatory Interpersonal Pleasure Scale, which measures anticipatory pleasure and consummatory pleasure with regard to social context (Gooding and Pflum, 2014). As such, it is critical to investigate reward-related behaviors to understand the mechanisms that drive anhedonia after ELA.

In behavioral tests explicitly targeting anhedonia, we observed the emergence of an ELA phenotype with aspects of anhedonia. Three-chamber social interaction is the most common test for reward processing of social interactions, where a reduction in direct social approach and time spent with a peer suggests an anhedonic phenotype (Der-Avakian and Markou, 2012; Scheggi et al., 2018; Trezza, V. et al., 2011). Our findings of significant reductions in direct social approach and average time spent with peer in ELA mice are similar to those previously reported in adolescent rats (Bolton et al., 2018a; Molet et al., 2016). Preweaned ELA rats had similar levels of social behavior, but after weaning adolescent ELA rats exhibited reduced social behavior (Rincon-Cortes and Sullivan, 2016). Sucrose preference also tests for anhedonia which is reduced after chronic stress (Papp et al., 1991). In our ELA mice sucrose preference was not significantly different from the controls over our three-day testing period, in contrast to published ELA rat studies (Bolton et al., 2018a; Molet et al., 2016). This may be a result of an elongated testing fourteen-day period in the rats compared to our abbreviated three-day testing period in mice. Our data suggests sociability deficits following ELA, which is a prominent symptom of anhedonia, as found in previous studies.

### 4.6 Relationship between social behavior and the NAcc

While exploring the possibility of interrelationships, we found that direct social approach was significantly correlated with regional FA, MD, AD, and RD of the NAcc. As such, decreased social approach in ELA mice was correlated with higher MD, AD, and RD and lower FA. This negative relationship suggests that sociability deficits are related to tissue-level diffusion changes observed in the NAcc of ELA mice. There is evidence to support the NAcc involvement in social behavior. One study of juvenile rats found that oxytocin within the nucleus accumbens supports a role in social novelty seeking behaviors (Smith et al., 2017). While another study found that µ-opioid receptors within the NAcc play a critical role in social reward (Trezza, Viviana et al., 2011). As such, alterations within the NAcc as reported by our diffusional changes likely underlie the social deficits observed in our ELA mice. Thus, that the DR-VTA-NAcc projection contributes to social reward processing and diffusion changes within the NAcc and may predict social deficits.

### 4.7 Limitations

While our findings are consistent with other ELA studies, our current study examined circuitry and reward-related behaviors only at a single time point. Of interest would be an examination of the temporal evolution that would better address when these circuits are modified after ELA. We also did not directly assess tissue-level cellularity which could be addressed using MRI T2 relaxation or cellular measures of histology to further illuminate the underlying changes we observed in both AD and RD. Finally, this study focused on males following ELA, therefore future studies will need to address sex differences following ELA.

### 4.8 Conclusion

We are the first to identify modifications of the DR-VTA-NAcc projection following ELA in male mice and to identify a unique ELA phenotype using a combination of DTI metrics, particularly AD and RD. Robust regional changes in RD, reward circuit connectivity, regional diffusion metrics, combined with along the tract AD together support the concept that ELA sculpts reward circuits as observed in adulthood. Furthermore, correlations identified relationships between direct social approach and the NAcc which supports our hypothesis that exposure to ELA leads to altered connectivity within the DR-VTA-NAcc projection. Although this was the first study to use diffusion metrics along the tract to pinpoint areas of altered organization within a projection after ELA, further investigation of the underlying cellular mechanisms and how these circuits develop during early-life brain development is necessary to understand how ELA perpetuates long term outcomes such as anhedonia.

## Acknowledgements

The authors would like to acknowledge the NIMH Silvio Conte Center, Conte Center on Brain Programming in Adolescent Vulnerabilities, University of California, Irvine. (2 P50 MH 096889 06A1, PI: T.Z. Baram). We would also like to acknowledge and thank Steven Granger for discussions and preliminary analyses for this study.

## Supplemental Figure Legends

**Supplemental Figure 1. FA and MD also exhibited brain wide regional changes after ELA.** Percent change in FA and MD of the ELA mice compared to controls also showed that many regions were altered after ELA. FA exhibited widespread decreases across almost all regions after ELA, whereas MD showed increases across majority of the regions following ELA. Pie charts show the distribution of regions with percent change that were increased versus decreased for the ELA group compared to controls.

**Supplemental Figure 2. Connectivity of the DR, VTA, and NAcc was altered in ELA mice.** The connectivity of each region highlights the alterations after ELA (*). The ELA group exhibited a stronger connectivity between the DR and the left VTA, and decreased connectivity between DR and the right NAcc (*). The left VTA connectivity was increased to the NAcc, but conversely decreased connectivity to the DR in the ELA group. In the ELA group, the left NAcc had increased connectivity to the left VTA but decreased connectivity to the left and right NAcc.

