## Supplementary figures and images for "Early Life Adversity in Male Mice Sculpts Reward Circuits"

### Supplemental Figure 1

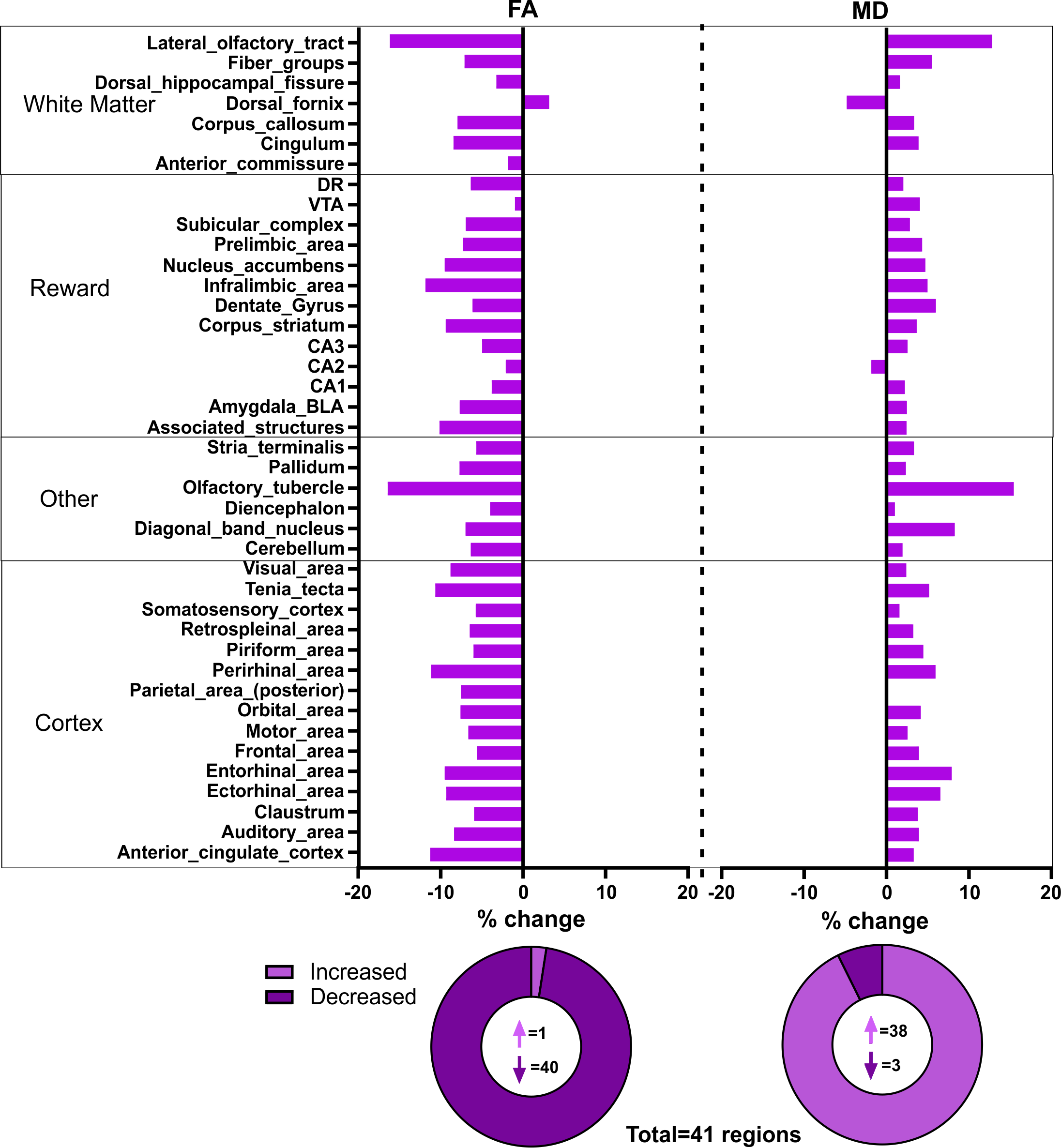

### Supplemental Figure 2

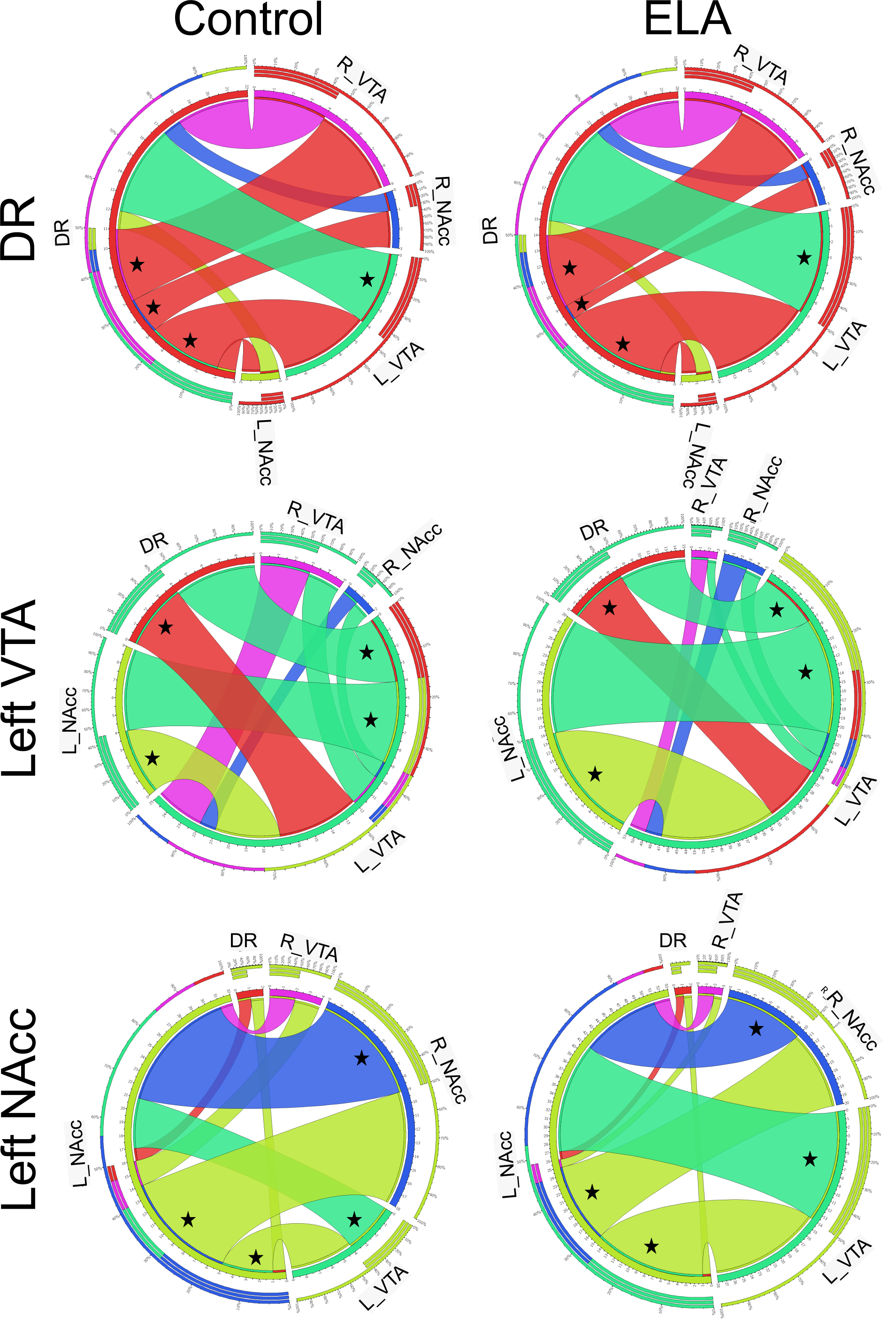
